# Parallel siRNA screens to identify kinase and phosphatase modulators of NF-κB activity following DNA damage

**DOI:** 10.1101/866061

**Authors:** Alexandros Sfikas, Peter Banks, Ling-I Su, George Schlossmacher, Neil D Perkins, Adrian I Yemm

**Affiliations:** Institute for Cell and Molecular Biosciences, Newcastle University, UK

## Abstract

DNA damage, such as that experienced by people undergoing chemotherapy, can directly activate NF-κB signalling which in turn can lead to resistance to genotoxic stress. NF-κB signalling is highly regulated by phosphorylation, but the enzymes required for these processes remain largely unknown. Identifying those enzymes responsible for regulating NF-κB activity may yield attractive targets for new clinical therapies, as well as provide the basis for better understanding of signalling network crosstalk. Here we present datasets from two independent RNAi screens using a stable NF-κB reporter U2OS cell line with the aim of identifying enzymes that alter NF-κB activity in response to DNA damage following etoposide and ionising radiation treatments. Although we observed high internal validity and specificity to NF-κB modulation within the screens, there was a striking dissimilarity between the results of the two different screens. These data therefore provide a cautionary lesson regarding the use of RNAi screening but also provide new candidates for kinase and phosphatase regulation of NF-κB activity in response to genotoxic stress.

## Background & Summary

The NF-κB family of transcription factors regulate a large and diverse set of target genes in response to many stimuli. Activation of NF-κB most commonly occurs through the canonical (classical) pathway. Typically, in this pathway, a heterodimer of the RelA(p65)/p50 NF-κB subunits becomes nuclear localised following degradation of its inhibitor, IκBα resulting from phosphorylation by the IκB kinase (IKK) complex. A wide range of stimuli can induce the canonical pathway, including many inflammatory cytokines and immune receptor pathways. However, in contrast to these signals emanating from the cell surface, the canonical NF-κB pathway can also be induced following nuclear DNA damage in a manner requiring the activity of the ataxia telangiectasia (ATM) kinase (Reviewed in ^1,2^).

Clinically, the response to exogenous DNA damage underpins genotoxic cancer treatment. Agents such as ionising radiation (IRR), and the topoisomerase inhibitor etoposide, induce DNA double strand breaks (DSB) to trigger cell death. In this context, NF-κB activation can contribute towards genotoxic therapy resistance in cancers through multiple routes ^3–6^.

Crosstalk between NF-κB and parallel signalling pathways provides a mechanism to regulate stimulus specific gene expression ^6,7^. Thus, identifying the kinases and phosphatases that mediate integration between NF-κB and other signalling networks following DNA damage may provide a novel strategy to target NF-κB signalling in cancer and combat genotoxic therapy resistance ^2^.

### Experimental Design

An unbiased approach to identifying kinases and phosphatases from parallel signalling pathways that modulate NF-κB signalling has not been previously explored, although several candidate based approaches have been reported ^8–10^. Here we aimed to identify novel kinases or phosphatases that regulate NF-κB activity following DNA damage by performing high throughput siRNA screens with 2 different inducers of DNA damage; etoposide and IRR.

We employed 2 independent siRNA screens to perform gene silencing of human kinases and phosphatases. Knockdown was performed in the human U2OS osteosarcoma cell line stably transfected with an NF-κB luciferase reporter plasmid. Following siRNA knockdown, cells were exposed to either etoposide or IRR and NF-κB activity was measured. Robust Z scoring was utilised to standardise results and identify potential siRNAs affecting NF-κB activity. Selected siRNAs were taken forward and a smaller repeat screen undertaken to determine reproducibility and specificity. Finally, some siRNAs were then used in small-scale experiments to further validate observations. The experimental flow path is depicted in Figure 1 and all experiments performed are listed in Table 1.

**Table 1:** Experiment List Table detailing experiments including experiment type, siRNA vendor, stimuli used in the experiment, the total number of siRNA included in each experiment, reporter cell lines used and affiliated data records with each study. * Data come from the same experiment. Unique Data Record as data handled differently to address different question. + Data Records include duplicate, extracted data from previous relevant data records

| Experiment Type | siRNA Vendor | Stimulus | No. siRNA | Cells | Affiliated Data Records |
| --- | --- | --- | --- | --- | --- |
| Initial Screen | Dharmacon | DMSO/Etoposide | 1124 | NFkB reporter U2-OS | 1, 2, 4, 8 |
|  |  | NON_IRR/IRR | 1124 | NFkB reporter U2-OS | 1, 2, 5, 9 |
|  | Sigma | DMSO/Etoposide | 719 | NFkB reporter U2-OS | 1, 3, 6, 10 |
|  |  | NON_IRR/IRR | 719 | NFkB reporter U2-OS | 1, 3, 7, 11 |
| Secondary Screen - "Reproducibility" | Dharmacon | DMSO/Etoposide | 74 | NFkB reporter U2-OS | 12*+ |
| Secondary Screen - "Specificity" | Dharmacon | DMSO/Etoposide | 74 | NFkB reporter U2-OS | 13* |
|  |  | DMOG | 74 | HRE U2-OS | 13* |
| Small Scale Validations | ONTARGET Plus | DMSO/Etoposide | 7 | NFkB reporter U2-OS | N/A |
|  |  | NON_IRR/IRR | 7 | NFkB reporter U2-OS | N/A |

**Figure 1:**
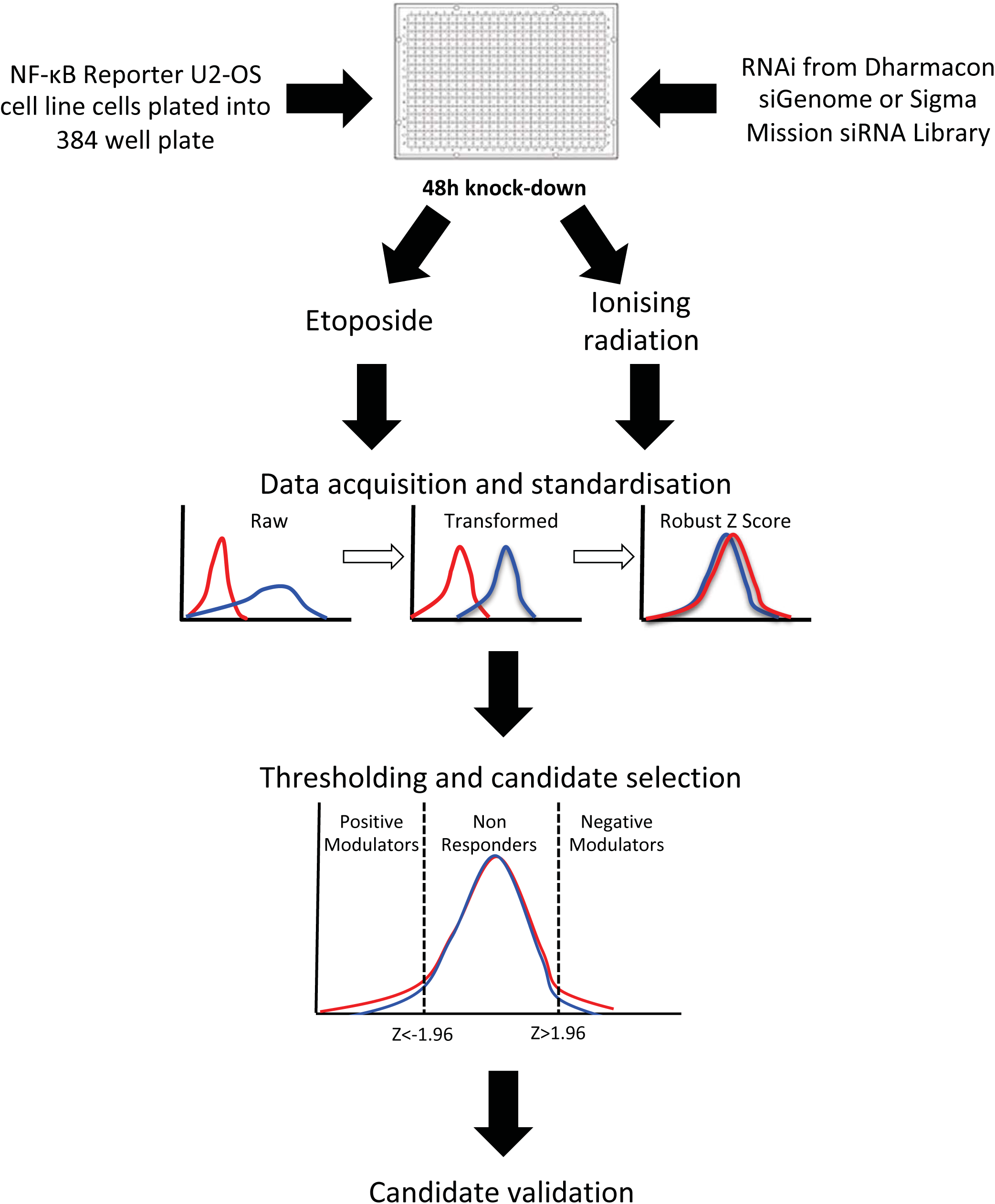
siRNA Screening Workflow Luciferase reporter cell lines are plated into 384 multiwell tissue culture plates at a density of 2000 cells/well and allowed to attach for 24h. Following culture, siRNA from either Dharmacon siGENOME Kinase Proteins Library or Sigma mission siRNA library are transfected at 10nM using Polyplus HTS transfection reagent in quadruplicate and with 4 biological repeats. Following 48h knockdown of target genes cells are either treated with 50μM etoposide (6h), 20Gy of Ionising Radiation for 6h or appropriate matched controls. Cells are then lysed in well and luciferase activity measured (BrightGlo, Promega; Omega POLARStar plate reader). Raw data are then processed and robust Z score calculated to determine effect of gene silencing.

We utilised 2 different commercially available siRNA screening platforms from Dharmacon and Sigma. The Dharmacon siGenome Protein Kinase Library consists of 1121 kinases and phosphatases and the Sigma Mission siRNA library consists of 715. Between the 2 screens there are 595 siRNAs in common, 526 unique to the Dharmacon screen and 120 unique to the Sigma screen. We predicted that there should be significant overlap between the siRNAs identified following etoposide and IRR treatments but that there would be unique stimulus specific modulators of NF-κB due to their different mechanisms of causing DNA damage.

We identified 111 positive and 143 negative regulators of NF-κB in total from the 2 different screens with a significant overlap in response to IRR and etoposide. Interestingly, both etoposide and IRR also had a number of unique modulators of NF-κB activity. A striking observation was the large disparity in the hits identified from siRNAs targeting the same genes between both vendors platforms, with as little as 3% overlap in response to IRR and 5% in response to etoposide. Further specificity and reproducibility studies revealed a high degree of internal reproducibility and specificity with the Dharmacon screen in our model system. However further studies using different siRNAs targeting a selection of these genes revealed poor validation of hits.

Overall, these data provide a candidate selection platform with which novel kinase and phosphatase regulators of NF-κB signalling can be pursued. Further studies may unveil mechanisms of integration between different signalling cascades as well as new candidates for pharmaceutical intervention in combination with genotoxic therapies for cancer.

However, these data sets also provide a cautionary tale with respect to screening platforms in which reagent choice and supplier can have a significant effect on candidate identification, emphasising that any candidates must be fully explored before any conclusions can be drawn.

## Methods

### Method 1 Cell lines and culture conditions

In order to quantify NF-κB activity in response to siRNA knockdown and stimulation we used the stably transfected pGL4.32[luc2P/ NF-κB-RE/Hygro (Promega) U2OS cell line described previously ^11^. The construct used to generate this stable cell line consists of 5 consecutive NF-κB consensus binding sequences to drive the expression of firefly luciferase and is used as a surrogate reporter for NF-κB transcriptional activity. This cell line was used in the primary screens and subsequent validation experiments. To confirm whether our observations were specific to NF-κB we used the reporter cell line HIF responsive element (HRE)-luciferase U2-OS described previously ^12,13^. The construct used to generate this stable cell line consists of the firefly luciferase expression driven by three HIF responsive elements from the Pgk-1 gene ^14^. These cells allowed us to confirm that any changes observed with siRNA modulation occurred specifically to NF-κB activity change and was not a general alteration to global transcriptional activity. With both cell lines, the use of a stable clonal cell line was to reduce variability as a result of transfection efficiency of the reporter constructs and to reduce population heterogeneity in baseline luciferase activity.

Both cell lines were maintained in Dulbecco’s modified Eagle’s medium (DMEM; Sigma-Aldrich) supplemented with 10% fetal bovine serum (FBS; Thermofisher), 2mM L-Glutamine (Lonza) and 200ug/ml G418 (Sigma-Aldrich) to maintain selection pressure on the stable cell lines.

### Method 2 Robotic handling

The Beckman Coulter Biomek FX was used to automate the dispensing of cells, transfection reagents, etoposide, and detection reagent in 384 well screening experiments.

### Method 3 siRNA screening

Using robotic handling, cells were plated into inner wells of white walled 384 multiwell plates at a density of 50000 cells/ml with 40 μl/well. Outer wells of plates were filled with cell suspension (but not included in any analyses) to reduce evaporation of experimental wells. Cells were then allowed to attach for 24h prior to transfection of siRNA. We utilised siRNA libraries from two different vendors for screening. Dharmacon siGENOME Protein Kinases Library (Purchase date: May 2012) and Sigma mission siRNA Library (Purchase date: May 2012). Table 2 outlines further details of siRNA libraries. Each initial screen was performed with 4 technical repeats with 4 independent biological replicates, secondary screens performed in quadruplicate. The Dharmacon platform also included RelA as a positive control. In all cases RelA knockdown was found to significantly reduce NF-κB activity. Any descriptions of candidates or hits in this manuscript do not include RelA as a component of the results due to its nature as a positive control.

**Table 2:**
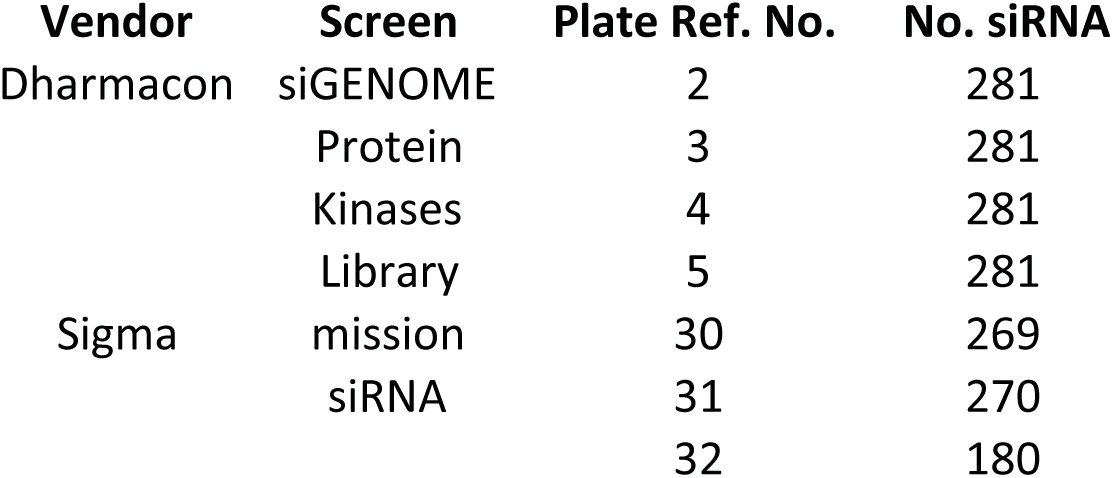
siRNA Libraries Overview

| Vendor | Screen | Plate Ref. No. | No. siRNA |
| --- | --- | --- | --- |
| Dharmacon | siGENOME | 2 | 281 |
|  | Protein | 3 | 281 |
|  | Kinases | 4 | 281 |
|  | Library | 5 | 281 |
| Sigma | mission | 30 | 269 |
|  | siRNA | 31 | 270 |
|  |  | 32 | 180 |

**Table 3:**
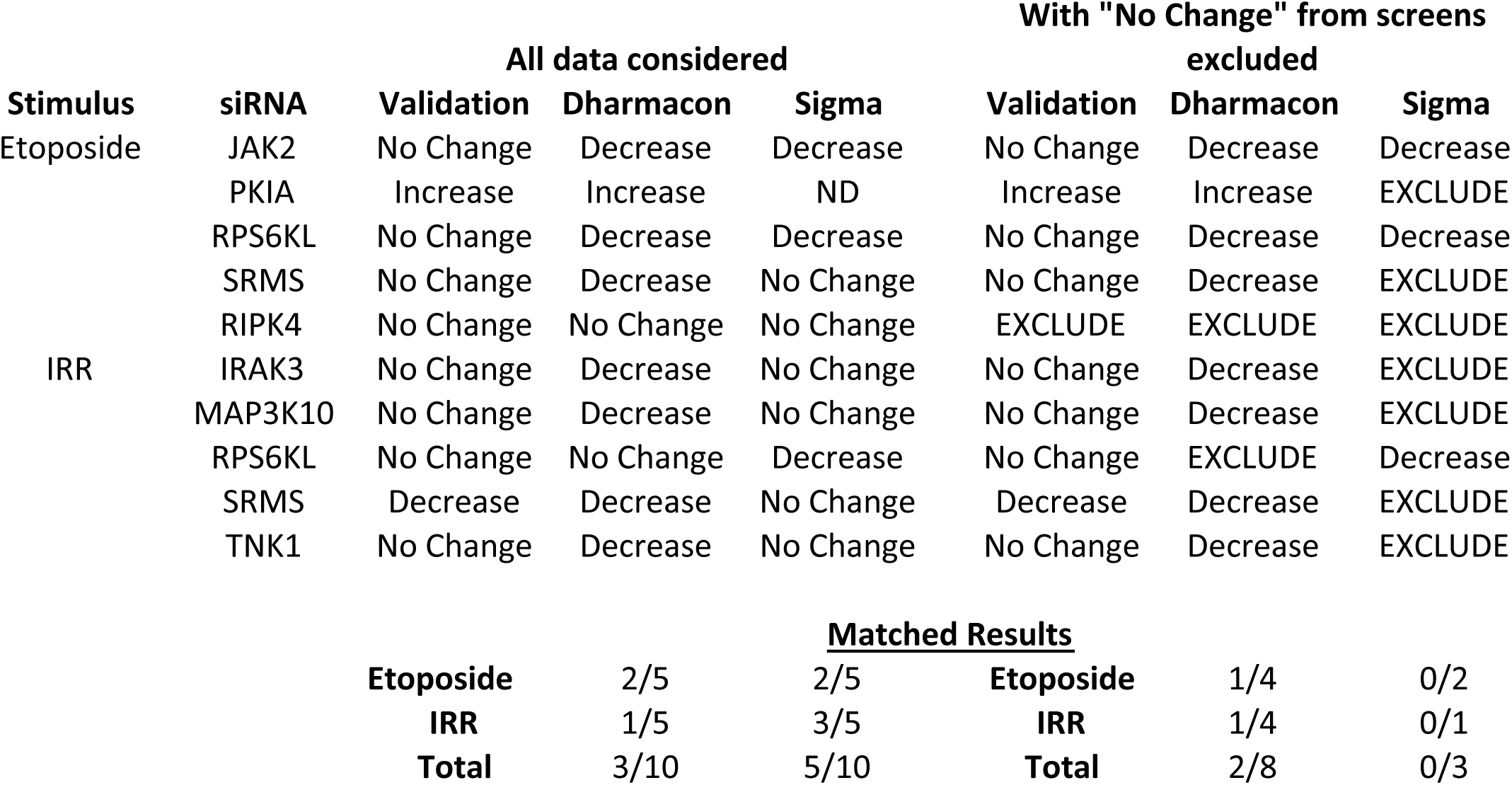
Summary table of validation results in comparison with screening results for selected siRNA. “No Change” highlights those siRNA that did not have an effect in the original screens. Given that most siRNA in these screens will have no effect on NFkB activity we suggest that this may be amore efficient way of considering validation efficiency. No Change = No significant change, Decrease = Significant decrease, Increase = Significant increase. For small scale validation results, significance determined by Student’s t-tests. For screening results robust Z score was used. An increase was considered >1.96, a decrease was considered <-1.96.

### Method 4 Transfection and stimulation procedure

Cells were transfected in well using Polyplus HTS (Polyplus) with a final concentration of 10nM siRNA. Cells were then incubated for 48h prior to stimulation. After 48h, transfection reagents were then aspirated and replaced with either etoposide or control media and handled as follows: 50μM Etoposide (E1383, Scientific Laboratory Supplies, UK) or DMSO (Sigma Aldrich), for 6 hours or 20Gy of ionising radiation (IRR) in growth media or handled in an identical manner without radiation and subsequently incubated for 6 hours. For HRE-luciferase U2OS experiments, cells were handled in an identical manner but stimulated with either DMSO or with the prolyl hydroxylase inhibitor dimethyloxaloylglycine (DMOG) at 500μM.

### Method 5 Luciferase activity quantitation

Following stimulation, cells were lysed in the wells using 35μl of BrightGlo (Promega, UK). Following lysis, data were acquired using a POLARstar Omega plate reader with stacker module (BMG Labtech). Luciferase activity was determined as relative light units (RLU) and exported to MS Excel for analysis. Luciferase activity in Specificity and Reproducibility screens (Stage 2 screens; see Figure 1, Table 1) were handled in an identical manner. For small scale validation experiments the Luciferase Assay System kit (Promega) was used as directed by manufacturers instructions and luminosity measured a using Lumat LB 9507 luminometer (Berthold Technologies).

### Method 6 Small scale validation experiments

Reporter U2OS cells were used in these experiments and maintained in an identical manner as above. Cells were plated into 6 well dishes (TPP) and grown to 30-50% confluency. The transfection protocol was as before; a 10nM final siRNA concentration, using Polyplus HTS interferin with the knockdown being performed for 48h prior to treatment. Scrambled control and selected siRNA were purchased from Dharmacon Ontarget plus SMARTpool or from Invitrogen: Scrambled: Dharmacon ONTarget plus Non-Targetting pool D-001810-10, RelA: 2 pooled custom siRNA; siRNA 1 5’ GCUGAUGUGCACCGACAAG, siRNA 2 5’ GCCCUAUCCCUUUACGUCA (Eurogentec). Dharmacon ONTarget plus SMARTpool: RIPK4 (54101) L-005308-00, JAK2 (3717) L-003146-00, SRMS (6725) L-005376-00, PKIA (5569) L-012321-00, IRAK3 (11213) L-004762-00, TNK1 (8711) L-003180-00, MAP3K10 (4294) L-003576-00. Invitrogen Stealth siRNA: RPS6KL (HSS130303), IKBKE (HSS114410). Experimental data were normalised to Scrambled siRNA controls and by control values for treatment. Normalised values were then subjected to Student’s t tests using Prism 6 (GraphPad).

### Method 7 siRNA screen analysis

For each repeat the following data handling steps were performed on a plate by plate basis. Initially, Log10 transformation of raw data was performed and the median and median absolute deviations (MAD) were calculated for each plate (Data Record 1). These values were then used to calculate Robust Z Scores for each siRNA in each biological repeat for each condition. A median absolute Z score of |1.96| was used to determine significant deviation from baseline response. Sign of the Z score represented significant increase or decrease in NF-κB activity. Candidate lists were then input into the online tool http://bioinformatics.psb.ugent.be/webtools/Venn/ to compare lists and draw Venn diagrams.

### Method 8 Reproducibility Assessment

To assess reproducibility “Rescreen” data (Table 1) were log10 transformed and median values determined. For comparison, data were extracted from relevant Data Records from “Original Screen” data (Table 1). The Log10 median values were subsequently ranked within each group, with lowest value =1 and ranked data were correlated. Spearman’s r correlations of ranked lists were calculated using Prism 6 (Graphpad).

### Method 9 Specificity assessment

To assess specificity “rescreen” data (Table 1) were log10 transformed and median values determined. NF-κB reporter data and HRE-luc data were ranked within each group, with the lowest value = 1. Then NF-κB reporter and HRE-luc values correlated using Spearman’s r in Prism 6 (Graphpad).

## Data Records

### Supplemental Data Record 1

**Data Record 1 = Summary Data**

Summary data for each plate used, including plate median and median absolute deviation values. These values are then used in Data Records 2-5 to allow for robust Z Score calculations for each siRNA studied under each condition (Method 7). The data record includes:

- siRNA_Screen – Supplier for siRNA screen
- No_siRNA – Total number of siRNA on plate
- Experiment – Stimulation conditions
- Plate_No – Experimental plate number
- Repeat_No – Independent biological repeat
- Plate_Median – Median value of all data within an experimental plate across biological repeats
- Plate_Log10_Median – Log10 transformation of Plate Median
- Plate_Log10_MAD – Median absolute deviation of plate

### Supplemental Data Record 2 – 3

**Data Record 2 = RefSeq Accession Number – Dharmacon Screen**

**Data Record 3 = RefSeq Accession Number – Sigma Screen**

RefSeq Accession Number for Dhamarcon (Data Record 2) and Sigma (Data Record 3) with associated gene name, plate number and well position. Each record is designated with:

- AcNumber – siRNA target gene accession number in RefSeq format
- Gene – Gene name associated with siRNA
- Plate – Plate reference number
- Row – Row number co-ordinate for siRNA within plate
- Column – Column number co-ordinate for siRNA within plate

### Supplemental Data Record 4 – 5

**Data Record 4 = Dharmacon Primary Screen Data – Etoposide**

**Data Record 5 = Dharmacon Primary Screen Data – Ionising Radiation**

Primary screen data for NF-κB reporter readout using the Dharmacon siRNA screen. Data Record 4 comprises data from Etoposide treatment and Control and Data Record 5 comprises data from Ionising Radiation treatment and Control. Each Data Record contains the raw data for each independent transfection for NF-κB activity (relative light units), Log10 transformations and Robust Z Score calculations for each repeat (Method 7). Each record is designated with:

- siRNA – Name of the gene the siRNA is targeting
- Plate – Source plate number from siRNA library
- C1-C4 – Raw RLU values for each repeat for control biological replicates
- T1-T4 – Raw RLU values for each repeat for stimulus biological replicates
- Log10C1-Log10C4 – Log10 transformation of RLU values for control biological replicates
- Log10T1-Log10T4 – Log10 transformation of RLU values for stimulus biological replicates
- ZC1-ZC4 – Robust Z score for each repeat for control biological replicates
- ZT1-ZT4 – Robust Z score for each repeat for stimulus biological replicates

### Supplemental Data Record 6 – 7

**Data Record 6 = Sigma Primary Screen Data – Etoposide**

**Data Record 7 = Sigma Primary Screen Data – Ionising Radiation**

Primary screen data for NF-κB reporter readout using the Sigma siRNA screen. Data Record 6 comprises data from Etoposide treatment and Control and Data Record 7 comprises data from Ionising Radiation treatment and Control. Each Data Record contains the raw data for each independent transfection for NF-κB activity (relative light units), Log10 transformations and Robust Z Score calculations for each repeat (Method 7). Each record is designated with:

- siRNA – Name of the gene the siRNA is targeting
- Plate – Source plate number from siRNA library
- C1-C4 (C1 – C6 for Data Record 7) – Raw RLU values for each repeat for control biological replicates
- T1-T4 (T1 – T6 for Data Record 7) – Raw RLU values for each repeat for stimulus biological replicates
- Log10C1-Log10C4 (Log10C1 – Log10C6 for Data Record 7)– Log10 transformation of RLU values for control biological replicates
- Log10T1-Log10T4 (Log10T1 – Log10T6 for Data Record 7)– Log10 transformation of RLU values for stimulus biological replicates
- ZC1-ZC4 (ZC1 – ZC6 for Data Record 7) – Robust Z score for each repeat for control biological replicates
- ZT1-ZT4 (ZT1 – ZT6 for Data Record 7) – Robust Z score for each repeat for stimulus biological replicates

### Supplemental Data Record 8 – 9

**Data Record 8 = Normalised Dharmacon Screen Data – Etoposide**

**Data Record 9 = Normalised Dharmacon Screen Data – Ionising Radiation**

Compiled normalised screen data calculated using the Robust Z Scores for the NF-κB reporter readout using the Dharmacon siRNA screen (Table 1). Data Record 8 comprises data from Etoposide treatment and Control and Data Record 9 comprises data from Ionising Radiation treatment and Control. The mean and standard deviation of calculated robust Z Scores (from Data Record 4 and 5) compiled for ease of interpretation. Each record is designated with:

- siRNA – Name of the gene the siRNA is targeting
- Plate – Source plate number from siRNA library
- C-Mean – Mean control Robust Z Score calculated from Data Records 4 – 5
- T-Mean – Mean stimulus Robust Z Score calculated from Data Records 4 – 5
- C-SD – Standard deviation of control Robust Z Score calculated from Data Records 4 – 5
- T-SD – Standard deviation of stimulus Robust Z Score calculated from Data Records 4 – 5

### Supplemental Data Record 10 – 11

**Data Record 10 = Normalised Sigma Screen Data – Etoposide**

**Data Record 11 = Normalised Sigma Screen Data – Ionising Radiation**

Compiled normalised screen data calculated using the Robust Z Scores for the NF-κB reporter readout using the Sigma siRNA screen (Table 1). Data Record 10 comprises data from Etoposide treatment and Control and Data Record 11 comprises data from Ionising Radiation treatment and Control. The mean and standard deviation of calculated robust Z Scores (from Data Record 6 and 7) compiled for ease of interpretation. Each record is designated with:

- siRNA – Name of the gene the siRNA is targeting
- Plate – Source plate number from siRNA library
- C-Mean – Mean control Robust Z Score calculated from Data Records 6 – 7
- T-Mean – Mean stimulus Robust Z Score calculated from Data Records 6 – 7
- C-SD – Standard deviation of control Robust Z Score calculated from Data Records 6 – 7
- T-SD – Standard deviation of stimulus Robust Z Score calculated from Data Records 6 – 7

### Supplemental Data Record 12

**Data Record 12 = Secondary Screen - Validation**

Compiled data from the secondary screen (Table 1) containing raw and log10 transformed RLU values. These data are from an experiment designed to replicate the results for selected siRNAs from the original screen (Method 8). These data were generated using NF-κB luciferase reporter U2OS cells only. The Data Record also contains a copy of data extracted from Data Records 4 – 7 (Designated “OriginalScreen”. Each record contains:

- siRNA – Name of the gene the siRNA is targeting
- Stimulus – Stimulus used in experiment. “Control” = untreated cells in the rescreen. “DMSO” = DMSO control from original screen. “Etoposide” = Etoposide treated cells from the original screen.
- Screen – Details which screen the data comes from. “OriginalScreen” = Data extracted from original screen (Data Record 4). “Rescreen” = Data generated in rescreen
- Raw1-Raw4 – Raw RLU values from experiment
- Log10-1 – Log10-4 – Log10 transformation of raw RLU values
- Log10Median – Median of Log10 transformed values
- Rank – Rank order of Log10Median values within each group (e.g. “Rescreen”, “Control”)

### Supplemental Data Record 13

**Data Record 13 = Secondary Screen - Specificity**

Compiled data from the secondary screen containing raw and log10 transformed RLU values. These data are from an experiment designed to compare selected siRNAs from the original screen using the NF-κB luciferase reporter U2OS cells (these data are duplicated from Data Record 12 Screen “Rescreen” for ease of analysis) and in HRE U2OS cells (Method 9). Each record contains:

- siRNA – Name of the gene the siRNA is targeting
- Stimulus – Stimulus used in experiment. “Control” = untreated cells, “DMOG” = DMOG treated HRE U2OS, “Etoposide” = Etoposide treated NF-κB reporter U2OS
- Cells – Reporter cell line data comes from (“HRE U2OS” = HIF1a Responsive Element U2OS, “NF-κB U2OS” = NF-κB reporter U2OS)
- Raw1-Raw4 – Raw RLU values from experiment
- Log10-1 – Log10-4 – Log10 transformation of raw RLU values
- Log10Median – Median of Log10 transformed values
- Rank – Rank order of Log10Median values within each group (e.g. “Rescreen”, “Control”)

## Technical Validation and Assessment of Data Quality

### Positional Effects

Method 10 was employed using data from Data Records 4-11 to determine whether any positional effects of the screening process could be identified. We confirmed that changes in NF-κB activity measured in these assays were not related to positional artefacts. This is apparent from the random distribution of luciferase activity across each of the plates. Figure 2 displays heatmap graphics of median log10 luciferase values.

**Figure 2:**
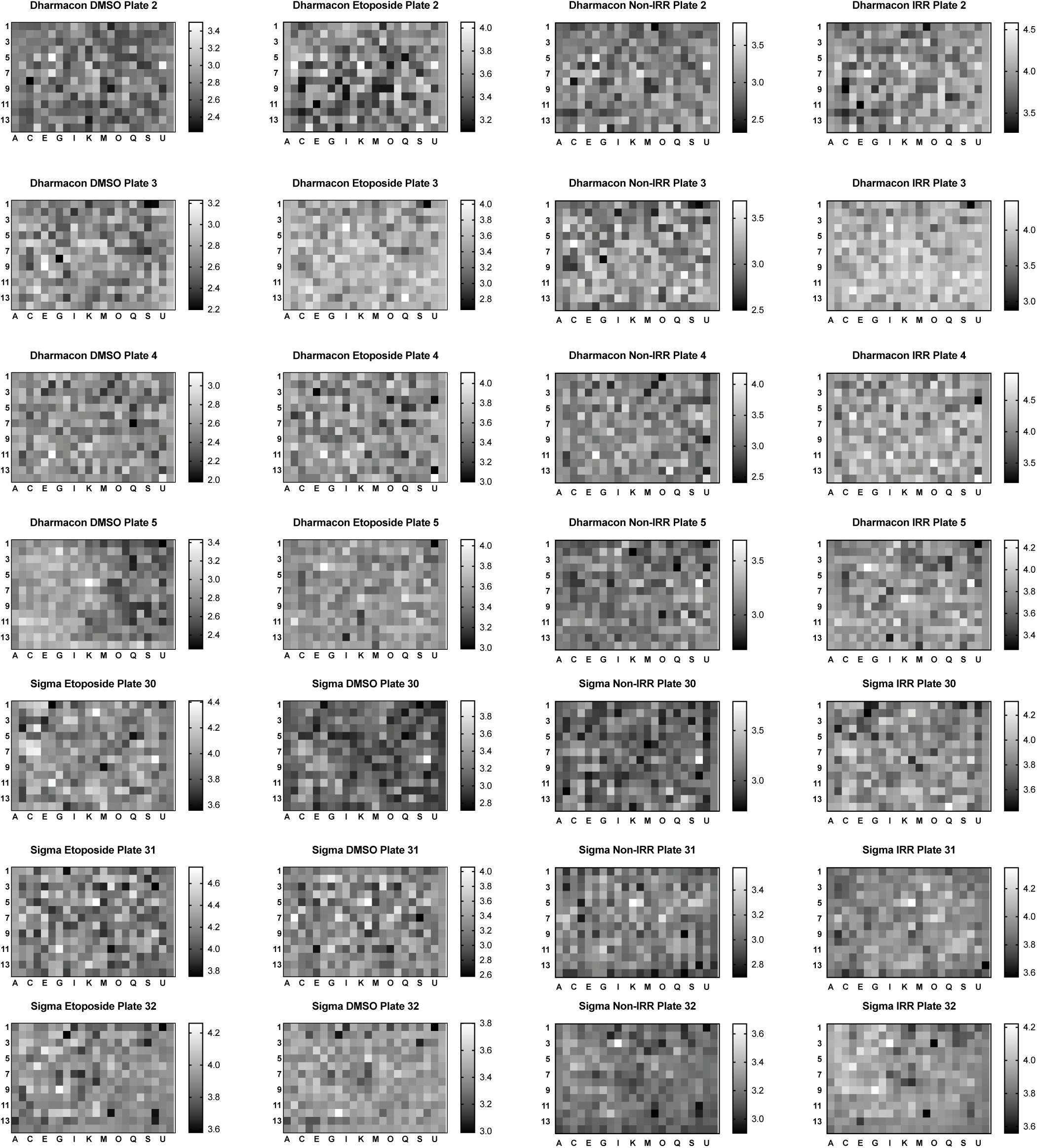
Screening procedure is not affected by positional artefact Average robust Z score for each well for each siRNA plate, for each condition studied, are presented as heat maps. The apparent random distribution of values across each plate confirms that positional artefacts caused by pipetting error or evaporation issues do not drive results.

### Transformation and Standardisation process is robust

Raw values were initially log transformed to improve the symmetry of the data (Method 7; Figure 3). Robust Z scoring was then used to allow for unbiased standardisation of scores in a manner resistant to outlier influence. In Figure 3 the data presented are from Data Record 4. These data are used as an example of the data handling process. This method allows for the identification of hits by selecting a Z score threshold (for example |1.96| is equivalent to 2 standard deviations from the mean) to determine siRNA that cause a significant change in NF-κB activity independently of treatment. Therefore, identifying those that cause a significant change from the ‘normal’ response either basally or following treatment. Importantly, Figure 3 also demonstrates that the transformation and standardisation process normalised any experimental variation across the different repeats.

**Figure 3:**
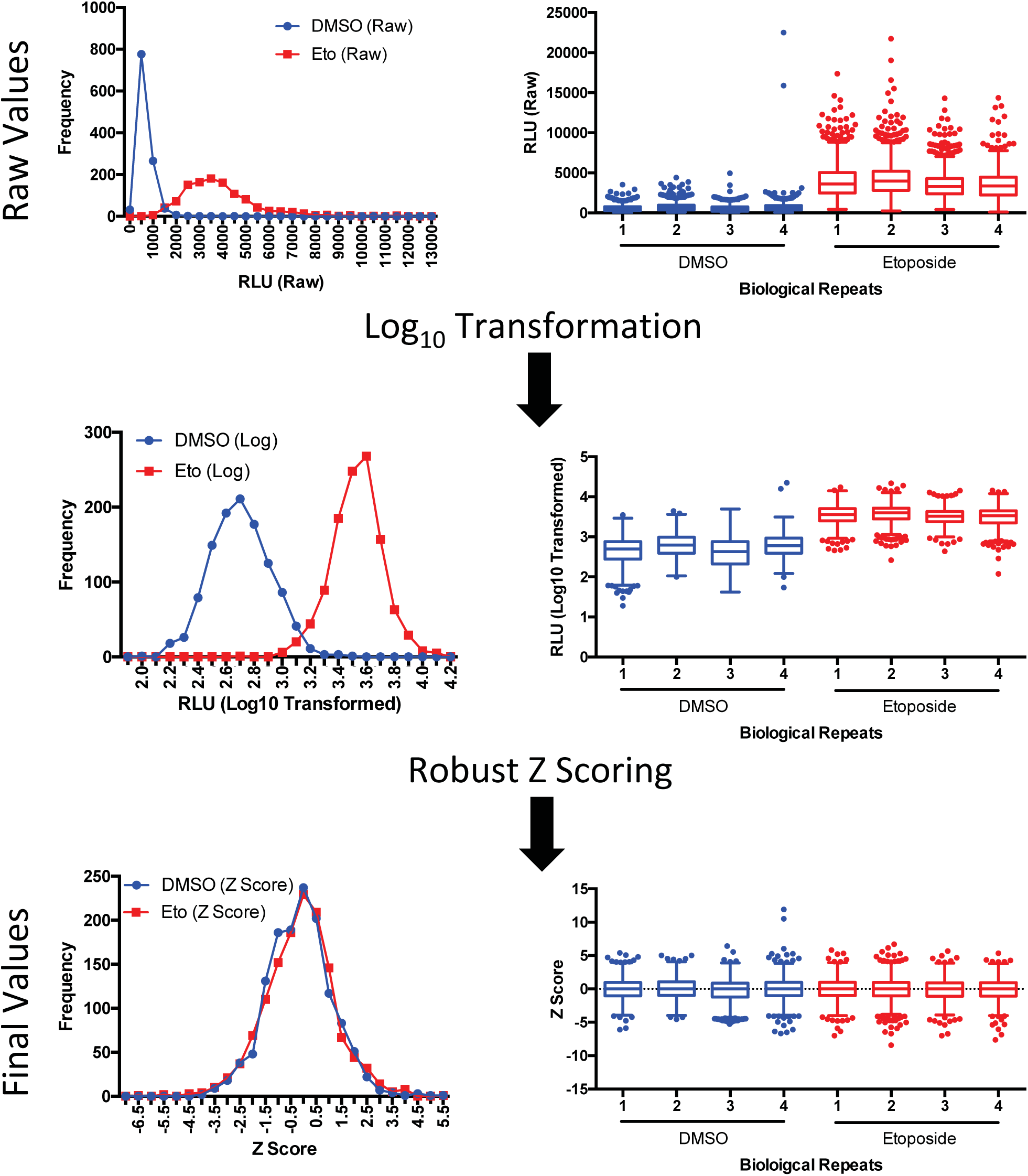
Graphical representation of transformation and standardisation process Frequency histographs and dot plots of data from Data Record 2 as an example of data processing steps. The processing of raw data is initially through Log10 transformation to make data more symmetrical. Following Log10 transformation robust Z Scoring is applied.

### The screening methodology applied is reproducible and specific

To determine the reproducibility of the screening process, a small panel of siRNA from the Dharmacon siRNA library were selected and used in a further repeat experiment (See “Secondary Screen” in Table 1; Method 1 and 8). The log transformed values were calculated and median determined for the new data set (Data Record 10) and median values from the original data set (Data Record 4) were extracted. These panels of siRNA were then ranked according to median response, with lowest value = 1. Ranked lists were then correlated using Spearman’s r to determine degree of correlation between the original screen and the rescreen. It was found that the rescreen correlated well with the original screen: Control vs DMSO r = 0.5125, p<0.0001; Etoposide vs Etoposide r = 0.5373, p < 0.0001. This provides a degree of confidence that using an siRNA high throughput screening approach to determine effects on NF-κB signalling is valid.

To provide confidence that the effects observed were specific to NF-κB signalling, the same panel of siRNA from the Dharmacon screen was also used with the HIF1a responsive element (HRE) reporter U2-OS cell line (Table 1) stimulated with DMOG (Method 1). As before, log transformed values were calculated and median response were ranked accordingly (Data Record 11). These ranked values were then correlated against the ranked values for NF-κB luciferase reporter U2-OS cells stimulated with etoposide and correlated. The failure to correlate between DMOG treated HRE cells and etoposide treated NF-κB luciferase reporter cells (DMOG vs Etoposide r = −0.04413, p = 0.7148) indicate that the changes in activity observed were specific to NF-kB activity changes.

Overall, these second screening studies confirm that siRNA screening in a high throughput manner is a reproducible and specific method for assaying for modulators of NF-kB.

#### Comparison of siRNA screening platforms

Due to the inherent off target effects common to siRNA based experiments we ran a comparative screen with an siRNA library from an alternative vendor. Following the analysis procedure, the lists of siRNAs having an effect on NF-κB activity from each experiment between conditions and between the Dharmacon and Sigma screens were compiled. Those siRNA with an absolute Z score of >|1.96| were input into http://bioinformatics.psb.ugent.be/webtools/Venn/ that compares lists and generates Venn diagrams.

Initially, we confirmed that there was significant overlap, as would be expected, between etoposide treated and IRR treated cells. Results from the Dharmacon screen identified that 42.75% of siRNAs either increased or decreased NF-κB activity (Figure 4A, Table 4) and 30.13% of candidates in the Sigma screen (Figure 4B, Table 4) following treatment with etoposide and IRR. Common negative regulators of NF-κB (i.e. those that result in increased NF-κB activity following gene silencing) made up 18.32% and 18.59% of the Dharmacon and Sigma screens respectively and positive regulators 24.43% and 11.54% (Table 4). These significant overlaps implied that the data are of high quality.

**Table 4:** Summary of overlap between etoposide and IRR within each screen * = significant change regardless of direction (i.e. an absolute Z score of |1.96|. An increase was considered significant if Z >1.96 and < −1.96 for decrease.

| Vendor | Stimulus | Change* | % Total | Increase | % Total | Decrease | % Total |
| --- | --- | --- | --- | --- | --- | --- | --- |
| Dharmacon<br>(262 siRNA) | Etoposide | 74 | 28.24 | 37 | 14.12 | 37 | 14.12 |
|  | IRR | 76 | 29.01 | 26 | 9.92 | 50 | 19.08 |
|  | Both | 112 | 42.75 | 48 | 18.32 | 64 | 24.43 |
| Sigma<br>(156 siRNA) | Etoposide | 71 | 45.51 | 36 | 23.08 | 35 | 22.44 |
|  | IRR | 38 | 24.36 | 14 | 8.97 | 24 | 15.38 |
|  | Both | 47 | 30.13 | 29 | 18.59 | 18 | 11.54 |

**Figure 4:**
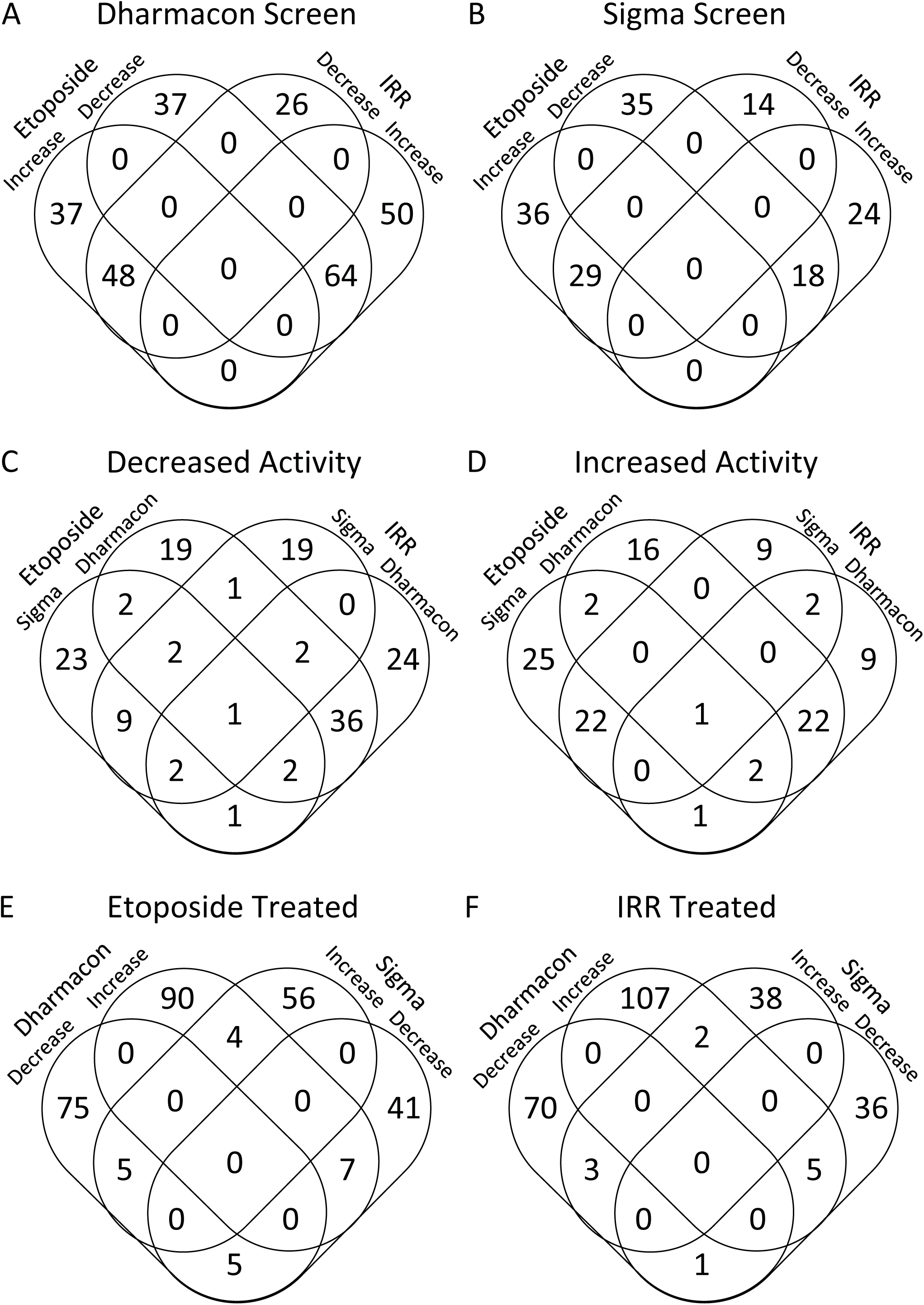
siRNA screens validate well internally but do not overlap between vendors. Venn diagrams are presented compiled from lists of candidates that were considered significantly increased (robust Z score >1.96) or significantly decreased (robust Z score < - 1.96) from the initial screen (Table1; Data Records 8-11) utilising NF-κB luciferase reporter U2-OS. A) Comparison of significantly different candidates (Z>|1.96|) between etoposide and IRR treated cells from Dharmacon screen (Data Record 8 & 9). B) Comparison of significantly different candidates (Z>|1.96|) between etoposide and IRR treated cells from Sigma Screen (Data Record 10 & 11). C) Comparison between etoposide and IRR treated cells from Dharmacon and Sigma screens that result in a significant decrease in activity (Z<-1.96). D) Comparison between etoposide and IRR treated cells from Dharmacon and Sigma screens that result in a significant increase in activity (Z>1.96). E) Comparison of etoposide treated cells that result in a significant increase or decrease in activity (Z>|1.96|) in either Dharmacon or Sigma screens. F) Comparison of IRR stimulated cells that result in a significant increase or decrease in activity (Z>|1.96|) in either Dharmacon or Sigma screens. In E and F, note the overlaps between increase and decrease groups in different screens. Data are also summarised in Tables 5–7.

After confirming high internal validity we also compared the screens from the two different vendors. Interestingly, there was a large discrepancy between the two screens, with very few common targets identified between the screens when performed under identical conditions. Comparing only those candidates from the list of siRNA that are shared between the two vendors only two siRNA were found to occur in all groups (Figure 4C & D; Table 5). This highlights the significant differences between the two screens, even though when considered separately they yield sensible and apparently reliable results.

**Table 5:** Summary of overlap of the screens between vendors * Sum of siRNA that exhibit some crossover between vendors that is not complete. E.g. 1 siRNA increased activity following etoposide treatment only in the Sigma screen and for IRR treatment only in the Dharmacon Screen.

| Vendor | Increase | Etoposide |  |  | Increase | IRR |  |  | Increase | Etoposide and IRR |  |  |
| --- | --- | --- | --- | --- | --- | --- | --- | --- | --- | --- | --- | --- |
|  |  | % of increased | Decrease | % of decreased |  | % of increased | Decrease | % of decreased |  | % of increased | Decrease | % of decreased |
| Dharmacon | 16 | 14.41 | 19 | 13.29 | 9 | 8.11 | 24 | 16.78 | 22 | 19.82 | 36 | 25.17 |
| Sigma | 25 | 22.52 | 23 | 16.08 | 9 | 8.11 | 19 | 13.29 | 22 | 19.82 | 9 | 6.29 |
| Shared | 2 | 1.8 | 2 | 1.4 | 2 | 1.8 |  |  | 1 | 0.9 | 1 | 0.7 |
| Partial* |  |  |  |  |  |  |  |  | 3 | 2.7 | 10 | 6.99 |
| Number of siRNA that increased NFkB activity across both vendors |  |  |  |  |  | 111 |  |  |  |  |  |  |
| Number of siRNA that decreased NFkB activity across both vendors |  |  |  |  |  | 143 |  |  |  |  |  |  |

Comparisons to this point had been undertaken considering only if a significant change is observed, not whether the change is an increase or decrease in activity. However, if the lists are recompiled to separately consider whether activity significantly increases or decreases then further discrepancies arise (Figure 4D & E; Table 6). When etoposide is considered, only 7.42% (21/283) of candidates were found to occur in both vendors but of these candidates 42.86% (9/21) display activity changes in opposite directions (i.e. some increase activity in one screen, but decrease activity in the other; Table 6). Similarly for IRR, only 4.20% (11/262) of candidates are found in both screens and of those 27.27% (3/11) drive NF-κB activity in opposite directions (Table 6). The results of these siRNA from different vendors to the same target gene driving NF-κB activity changes in opposite directions (depending upon the source of the siRNA) are striking and highly concerning observations.

**Table 6:** Summary of divergent results between vendors * = significant change regardless of direction (i.e. an absolute Z score of |1.96|. An increase was considered significant if Z >1.96 and < −1.96 for decrease.

| Stimulus | Vendor | Change* | % Total | Increase | % Total | Decrease | % Total | Opposite | % of Change | % Total |
| --- | --- | --- | --- | --- | --- | --- | --- | --- | --- | --- |
| Etoposide<br>(283 siRNA) | Dharmacon | 165 | 58.3 | 75 | 26.5 | 90 | 31.8 |  |  |  |
|  | Sigma | 97 | 34.28 | 56 | 19.79 | 41 | 14.49 |  |  |  |
|  | Both | 21 | 7.42 | 5 | 1.77 | 7 | 2.47 | 9 | 42.86 | 3.18 |
| IRR<br>(262 siRNA) | Dharmacon | 177 | 67.56 | 70 | 26.72 | 107 | 40.84 |  |  |  |
|  | Sigma | 74 | 28.24 | 38 | 14.5 | 36 | 13.74 |  |  |  |
|  | Both | 11 | 4.2 | 3 | 1.15 | 5 | 1.91 | 3 | 27.27 | 1.15 |

\* = significant change regardless of direction (i.e. an absolute Z score of |1.96|. An increase was considered significant if Z >1.96 and <1.96 for decrease.

The poor overlap between the two vendors is unexpected due to the validity testing performed on the Dharmacon screen (Data Record 10) and the high internal validity observations (Figure 4A & B; Table 4). It is likely that both sets of reagents are performing as described, but the off target effects inherent in RNA interference likely results in different perturbations on a signalling system as responsive as NF-κB. Also, it is likely that the different siRNA used for each target across the two vendors differ in efficiency that may also yield differential effects.

### The RNAi platforms replicate most known modulators of NF-κB activity

NF-κB mediated response to DNA damage has been well studied, and as a result there are several known kinases that have been shown to be involved in modulating NF-κB signalling in response to DNA damage. These candidates have been compiled in Table 7. The Dharmacon screen had a similar expected response to the literature for 6/8 candidates in response to etoposide and 5/8 in response to IRR. Interestingly, there was no effect seen following knockdown of IKBKG (NEMO), a known required enzyme for NF-κB activity in response to DNA damage ^2^. We speculate that this is likely due to inefficient knockdown of NEMO in these samples. It should also be noted that when these candidates did not match the literature, no effect was observed in the screen. The Sigma screen replicated the literature in 4/7 cases for both etoposide and IRR treatments. One notable result was that CHUK (IKKα) had no effect on NF-κB signalling. Again, we can only surmise that these siRNA were ineffectual in the Sigma screen. Unfortunately IKBKG (NEMO) did not form part of the Sigma siRNA platform.

**Table 7:**
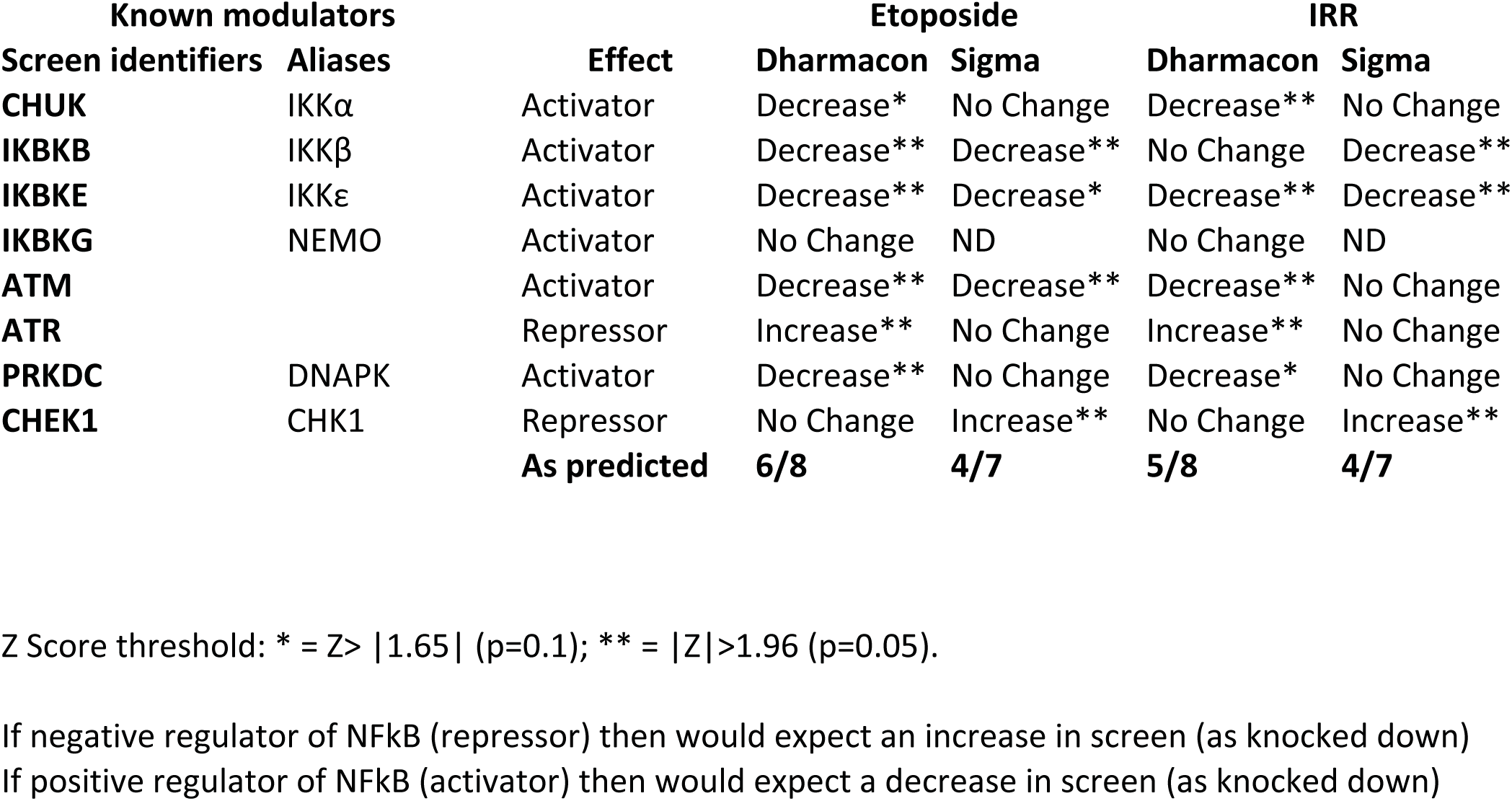
Summary table of known modulators of NF-κB and results in screens. A known repressor of NF-κB (i.e. normally reduces NF-κB activity) when knocked down by RNAi would result in an increase in NF-κB activity. Conversely, a known activator of NF-κB, when knocked down would result in a decrease in NF-κB activity. Robust Z Score threshold: * = Z> |1.65| (p=0.1); ** = |Z|>1.96 (p=0.05).

| Known modulators |  |  | Etoposide |  | IRR |  |
| --- | --- | --- | --- | --- | --- | --- |
| Screen identifiers | Aliases | Effect | Dharmacon | Sigma | Dharmacon | Sigma |
| CHUK | IKKα | Activator | Decrease* | No Change | Decrease** | No Change |
| IKBKB | IKKβ | Activator | Decrease** | Decrease** | No Change | Decrease** |
| IKBKE | IKKε | Activator | Decrease** | Decrease* | Decrease** | Decrease** |
| IKBKG | NEMO | Activator | No Change | ND | No Change | ND |
| ATM |  | Activator | Decrease** | Decrease** | Decrease** | No Change |
| ATR |  | Repressor | Increase** | No Change | Increase** | No Change |
| PRKDC | DNAPK | Activator | Decrease** | No Change | Decrease* | No Change |
| CHEK1 | CHK1 | Repressor | No Change | Increase** | No Change | Increase** |
| As predicted |  |  | 6/8 | 4/7 | 5/8 | 4/7 |

Z Score threshold: \* = $Z > |1.65|$ (p=0.1); \*\* = $|Z| > 1.96$ (p=0.05).
If negative regulator of NFkB (repressor) then would expect an increase in screen (as knocked down) If positive regulator of NFkB (activator) then would expect a decrease in screen (as knocked down)

Overall, the comparison with known candidates provides a fair degree of confidence in the suitability of RNAi screening for NF-κB activity modulators. It is interesting to note, that the “failed” observations of one screen were correct in the other screen. This further implies a failure of siRNA providing the negative result rather than those targets not having an effect. Overall, however, these observations further reinforce the need for further validation with RNAi screens as false negatives, due to siRNA failure for example, cannot be readily identified.

### Small Scale Validation

To further validate the screen a series of smaller scale experiments were conducted using different siRNA from those in either screen. OnTarget plus siRNAs (Dharmacon), similar to the Dharmacon and Sigma screens, contain a pool of four different siRNA targeting a single gene target (Method 6), however, they are designed to have fewer off target effects by utilising Dharmacon’s patented dual-strand modification design. These experiments used a selection of different siRNA (Table 1, Method 6) for validation following etoposide treatment (Figure 5A) or following IRR stimulation (Figure 5B).

**Figure 5:**
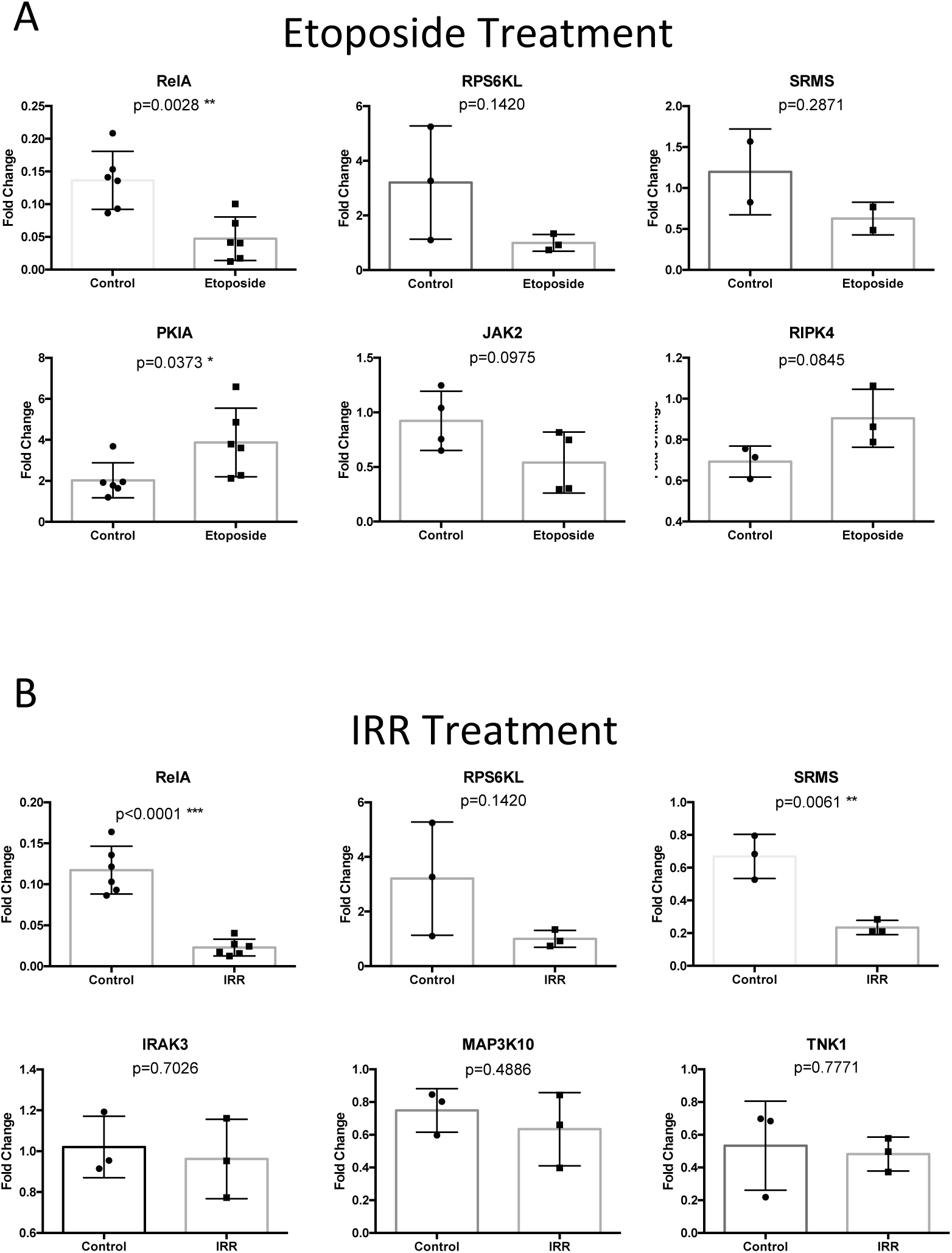
Small scale validation does not support observations from siRNA screens A) 7 siRNA were selected to validate observations seen in screens following etoposide stimulation. B) 7 siRNA selected to validate observations seen in screens following IRR treatment. Data are luciferase activity measurements normalised to Scrambled controls and expressed as Fold Change. Statistical evaluation performed using Student’s t-test, p values are reported for each experiment. Data points are individual experiments. Data are also summarised in Table 3.

In summary, within the etoposide experiments, 2/5 of Dharmacon siRNAs were validated, and 2/5 of the Sigma siRNAs were validated, with an overall average of 2/5 (Table 3). Similarly, the IRR experiments did not validate well, with small-scale experiments validating 1/5 of Dharmacon results and 3/5 of Sigma results, giving an average of 2/5 (Table 3). Combined, the Dharmacon siRNA screen results were validated with 3/10 of siRNAs while 5/10 of siRNAs were validated for the Sigma screen (Table 3). However, perhaps a more representative consideration of validation results is to exclude the “No Change” results obtained in the screen. Results considered “No Change” are those that are found to not alter NF-κB activity at all in the screening results. Given that the majority of siRNA were likely to have no significant effect, we predict that by removing these “No Change” results, we may get a more true reflection of the validation statistics. Summary statistics were recalculated (Table 3) and it was found that now the Dharmacon screen fell to only 1/4 of probable hits being validated while the Sigma screen fell to 0% validated. Therefore, across the two screens, on average 1/6 of etoposide related changes and 1/5 of IRR related changes were validated, giving an overall average of 2/11 of observations validated (Table 3).

These limited levels of validation performed using different siRNA to the screens provide cautionary evidence regarding the ability to draw robust conclusions from high throughput RNA interference screens and the inherent variability in such screens resulting from off-target effects.

#### Overall summary of data quality

These data, by our measures, show confidence in the validity of the screening process and that these data sets can yield useful, informative observations on NF-κB signalling in response to DNA damage. However, we believe these data strongly reinforce the need for further validation of any putative targets prior to any conclusions being drawn, particularly from a single set of reagents from a single supplier. This opinion is strongly supported by our observations that the siRNA from two separate vendors yield significantly different results. Caution must be applied to any high throughput study and limitations always borne in mind.

## Acknowledgements

Funded in part by Cancer Research UK grants C1443/A12750 and C1443/A22095 and BBSRC grant BB/L006480

## Author contributions

AS: Performed initial RNAi screens and designed experiments

PB: Performed initial RNAi screens and designed experiments

LS: Performed small scale validation screens

GS: Performed small scale validation screens

NDP: Designed experiments, interpreted results and wrote manuscript

AIY: Designed experiments, analysed data, interpreted results and wrote manuscript

